# Cell-Line Annotation on Europe PubMed Central

**DOI:** 10.1101/011700

## Abstract

A cell line is a cell culture developed from a single cell and therefore consisting of cells with a uniform genetic make-up. A cell line has an important role as a research resource such as organisms, antibodies, constructs, knockdown reagents, etc. Unique identification of cell lines in the biomedical literature is important for the *reproducibility* of science. As data citation, resource citation is also important for resource re-use.

In this paper, we mention the challenges of identifying cell lines and describe a system for cell line annotation with perluminary results.

## 1 Introduction

### 1.1 Why do we tag cell lines?

A cell line is a cell culture developed from a single cell and therefore consisting of cells with a uniform genetic make-up. A cell line has an important role as a research resource such as organisms, antibodies, constructs, knockdown reagents, etc. Unique identification of cell lines in the biomedical literature is important for the *reproducibility* of science [7]. As data citation, resource citation is also important for resource re-use [1]. Identifying cell lines is a non-trivial problem with the following challenges and difficulties:

- A significant number of cell line names consist of only numbers.
- A significant number of cell line names consist of less than 4 letters.
- Cell line names often look similar with gene/protein names.
- Cell line names sometime look similar with person names.

### 1.2 Linking Europe PMC articles to cell lines

Europe PubMed Central is a database of life science research articles and abstracts, including PubMed (http://europepmc.org) [4]. One of main services on Europe PMC is to link full-text articles to biological data sets or databases by two methods:

- Named Entity Recognition
- Accession Number Mining [2]

Combined with other features on Europe PMC, cell line annotation can be useful. For example, give me all articles where cell line X is mentioned only in Methods section.

In this report, we describe our work on linking articles to research resources using our cell line tagger and section tagger.

## 2 A large-scale annotation and analysis pipeline

Recently, we have developed a system which can generate a dictionary from a given ontology or terminological resource, and performs a large scale analysis of dictionary usages. The system mainly consists of three modules: 1) dictionary building module, 2) semantic tagging module, and 3) analysis and report generation module. Figure 1 shows an diagram of the system architecture.

**Fig. 1:**
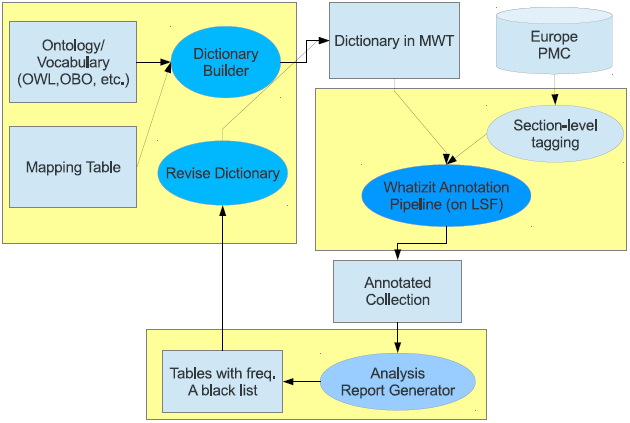
A pipeline for a large scale annotation and analysis given an ontology.

### 2.1 Dictionary builder

Dictionary builder is a module that generates a dictionary (in mwt format used in Whatizit), given an ontology, controlled vocabulary, or any other terminological resources. A number of input formats supported are as follows:

- Web Ontology Language (OWL) Recently, there has been a trend to develop ontologies in OWL recommended by W3C^1^. This module can generate dictionaries from ontologies in Web Ontology Language (OWL) using SPARQL Query Language for RDF.
- Open Biomedical Ontologies (OBO)
- Swiss-Prot style format

Besides the given ontology, additional information is required for building a dictionary as follows:

- A mapping table between an ontology and a dictionary (as in Table 1), and *is-a* relationship if any exists.
- A list of filtering rules. For example,
  Remove a term of which the length is less than N.
  Remove a term that has only digits.
- A list of regular expressions (e.g., for accession numbers)

**Tab. 1:**
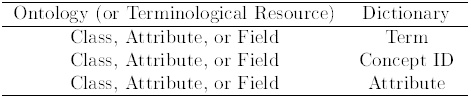
A mapping table between an ontology and a dictionary. In general, one concept ID has one or more than one terms.

| Ontology (or Terminological Resource) | Dictionary |
| --- | --- |
| Class, Attribute, or Field | Term |
| Class, Attribute, or Field | Concept ID |
| Class, Attribute, or Field | Attribute |

### 2.2 Semantic tagging engine

This module consists of a list of taggers based on java finite automata [5] and MALLET (MAchine Learning for LangugE Tookit^2^): Given a dictionary (generated in the previous step) in mwt^3^ format, this annotation pipeline can annotate documents in a dictionary-based approach. We can plug in different semantic taggers running on Whatizit server.

### 2.3 Analysis and summary report generation module

A large scale annotation analysis module based on Hadoop, Pig Latin (for counting), and R language (for visualization). One application of this analysis module is to help users with building a customized list of stop words and revising a dictionary based on summary report.

One method to evaluate the importance of a term is to us its frequency [3]. This frequency information can be used to find a list of stop words for domain-specific applications instead of using a default list of stop words.

## 3 Results

### 3.1 Cellosaurus-based cell-line dictionary

We have performed a preliminary analysis on Cellosarus and Cell Line Ontology (CLO). In this analysis, we have found that terms in Cellosarus are more often matched on free texts from biomedical corpora, suggesting the usage as a dictionary. On the other hand, CLO terms are more conceptual and less matched.

Based on this, we have chosen Cellosaurus which is a controlled vocabulary of cell lines developed by Swiss Institute of Bioinformatics (SIB).

With the dictionary building module (mentioned in Section 2.1), we generated a cell-line dictionary for named entity recognition as follows:

- In a mapping table, we mapped ID (IDentification) and SY (SYnonyms) fields to terms, and AC (ACcession number) field to concept IDs.
- Following is a list of filtering/ *transformation* rules:
  Only terms with more than 3 letter terms are used to build dictionaries.
  Less than four letters + ’cell’ (as the following constraint word)
  Only numbers + cell
- Following is a list of terms added into our blacklist, provided by domain experts:
  Cancer, Center, Grey, Spindle, Chance, Patches, Bones, Horse, TIME, Set-2, Renal carcinoma, Badger, Chew, Moose, Marry, Scout, COST, Pin-wheel, Giant cell tumor, Fetch, Mint, CHOP, Ears, Jersey, Chase, Chief, Flip, Guard, Junior, Stripes, Squirrel, Typhoon, Sage, Had-1, Speedy, Thyme, WISH, Kin-, Tackle, Pepper, Taurus, WART, Speckles, Soccer, Buttons, Gemini, Bing (47 terms)

Table 2 shows statistics on the dictionary built by this module and Figure 2 shows an example of the dictionary generated based on these rules.

**Tab. 2:** About Cellosaurus-based (version 8.0) cell-line dictionary.

|  | Before filtering | After filtering |
| --- | --- | --- |
| # Concepts | 23,115 | 23,107 |
| # Terms | 39,089 | 39,042 |

**Fig. 2:**
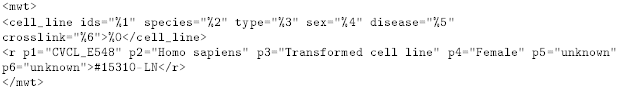
Dictionary example

### 3.2 Annotation results on Open Access (OA) PMC collection

For a large scale analysis, first we applied our section tagger to 633,174 OA full-text articles from Europe PubMed Central collection, and annotated these section-labeled articles. The rational for using section-labeled articles is to find different usages of cell lines over different sections such as Introduction, Methods, Results, and Discussion [6]. Then, we applied our gene/protein name tagger in order to reduce confusions between gene/protein and cell line names.

After the annotation we performed three different analyses: concept-wise, term-wise, and article-wise in order to find different aspects of ontology usages.

In term-wise analysis, each term was counted separately. Table 3 shows the results. In this table, we can see section-specific false positive cases. For example, there are some false positive terms specific in Methods section such as Fisher and Madison. Different sections have different false positive categories suggesting needs for a section-dependent blacklist.

**Tab. 3:** Top 30 most frequent terms (collection frequency). A number of chunks for each section is as follows: 57,291,268 for Introduction, 98,112,562 for Methods, 156,007,208 for Results, and 83,851,662 for Discussion.

| Introduction |  | Methods |  | Results |  | Discussion |  |
| --- | --- | --- | --- | --- | --- | --- | --- |
| HeLa | 3068 | Fisher | 28966 | HeLa | 52563 | HeLa | 7662 |
| TLR4 | 1934 | *Madison* | 23793 | MCF-7 | 21262 | MCF-7 | 5984 |
| MCF-7 | 1778 | HeLa | 21689 | HEK293 | 17357 | TLR4 | 4173 |
| Fisher | 1499 | 293T | 9073 | HepG2 | 14381 | HepG2 | 3695 |
| TLR2 | 1353 | HEK293 | 8671 | 293T | 12425 | TLR2 | 3380 |
| HepG2 | 1218 | MCF-7 | 6663 | HCT116 | 11396 | PC12 | 3200 |
| PC12 | 1205 | BL21 | 5722 | MCF7 | 10412 | LNCaP | 2499 |
| *Jensen* | 1191 | HepG2 | 5299 | LNCaP | 9574 | MDCK | 2341 |
| *Hughes* | 1156 | Vero | 5202 | PC12 | 8082 | HEK293 | 2315 |
| *Murphy* | 1077 | MDCK | 4515 | MDCK | 7872 | MCF7 | 2044 |
| *Becker* | 841 | HEK293T | 4497 | C2C12 | 7622 | 3T3-L1 | 1773 |
| *Kobayashi* | 739 | *Focus* | 3981 | HEK293T | 7324 | C2C12 | 1701 |
| LNCaP | 737 | HCT116 | 3246 | U2OS | 6589 | PC-3 | 1682 |
| Cole | 710 | MCF7 | 3000 | Fisher | 6543 | HCT116 | 1653 |
| HEK293 | 703 | C2C12 | 2976 | SH-SY5Y | 6393 | Fisher | 1481 |
| LC-MS | 681 | PC12 | 2923 | TLR4 | 5738 | SH-SY5Y | 1391 |
| C2C12 | 670 | LNCaP | 2721 | PC-3 | 5408 | Jensen | 1306 |
| MDCK | 668 | LC-MS | 2710 | Vero | 5222 | SP cells | 1245 |
| 3T3-L1 | 663 | Fuji | 2657 | NIH3T3 | 5201 | HT-29 | 1169 |
| Fang | 661 | COS-7 | 2495 | TLR2 | 4877 | Murphy | 1164 |
| Vogel | 570 | SH-SY5Y | 2434 | 3T3-L1 | 4642 | Hughes | 1077 |
| MCF7 | 540 | F4/80 | 2431 | HaCaT | 4498 | NIH3T3 | 1026 |
| Otto | 499 | U2OS | 2251 | COS-7 | 4376 | HT29 | 1024 |
| PC-3 | 488 | NIH3T3 | 2246 | SW480 | 4333 | Vero | 1016 |
| SP cells | 479 | 293 cells | 2207 | HT29 | 4320 | HaCaT | 972 |
| Peer | 473 | 3T3-L1 | 1919 | DU145 | 4197 | Kobayashi | 943 |
| Focus | 469 | HEK 293 | 1849 | RAW264.7 | 3846 | DU145 | 932 |
| SH-SY5Y | 463 | RAW 264.7 | 1844 | HT-29 | 3759 | 293T | 883 |
| DT40 | 452 | RAW264.7 | 1813 | H1299 | 3676 | RAW264.7 | 847 |
| HCT116 | 443 | HEK-293 | 1790 | T47D | 3661 | Becker | 804 |

- Kobayashi in Introduction vs Kobayashi in Methods
- Promega Corporation, *Madison*, WI, USA.

Based on the results above, we removed term *Focus*, which is often used as “Focus group” in Methods sections.

HeLA was not the most frequent term in Methods section because of two false positive terms (i.e., Fisher and Madison).

Concept-wise analysis: All terms (synonyms, orthographic variants, etc) belonging to one concept were considered as the same. Table 4 shows 15 most frequent concepts.

**Tab. 4:** Top 15 most frequent concept IDs. In general, one concept ID has one or more than one terms. For example, CVCL_0063 has HEK293T, HEK 293T, 293T, and any other terms.

| Introduction |  | Methods |  | Results |  | Discussion |  |
| --- | --- | --- | --- | --- | --- | --- | --- |
| CVCL_0030 | 3072 | CVCL_E017 | 28966 | CVCL_0030 | 52629 | CVCL_0031 | 8056 |
| CVCL_0031 | 2318 | CVCL_H602 | 23835 | CVCL_0031 | 31782 | CVCL_0030 | 7679 |
| CVCL_F956 | 1934 | CVCL_0030 | 21712 | CVCL_0045 | 26996 | CVCL_F956 | 4173 |
| CVCL_E017 | 1499 | CVCL_0063 | 16246 | CVCL_0063 | 22922 | CVCL_0027 | 3805 |
| CVCL_5600 | 1353 | CVCL_0045 | 14600 | CVCL_0027 | 14677 | CVCL_0045 | 3713 |
| CVCL_0481 | 1278 | CVCL_0031 | 9697 | CVCL_0291 | 13772 | CVCL_5600 | 3380 |
| CVCL_0027 | 1268 | CVCL_M639 | 6014 | CVCL_0594 | 10757 | CVCL_0481 | 3348 |
| CVCL_3531 | 1191 | CVCL_0027 | 5478 | CVCL_0395 | 9900 | CVCL_0395 | 2576 |
| CVCL_0045 | 1158 | CVCL_0059 | 5370 | CVCL_0035 | 8513 | CVCL_0035 | 2357 |
| CVCL_L357 | 1156 | CVCL_0594 | 5295 | CVCL_0481 | 8446 | CVCL_0422 | 2341 |
| CVCL_3549 | 1077 | CVCL_0422 | 4521 | CVCL_0320 | 8079 | CVCL_0320 | 2193 |
| CVCL_0594 | 877 | CVCL_7955 | 4130 | CVCL_0493 | 8022 | CVCL_0594 | 2175 |
| CVCL_1093 | 841 | CVCL_0224 | 4123 | CVCL_0422 | 7878 | CVCL_0291 | 2147 |
| CVCL_0395 | 766 | CVCL_0291 | 4113 | CVCL_0188 | 7657 | CVCL_0123 | 1902 |
| CVCL_J354 | 739 | CVCL_0493 | 3901 | CVCL_0042 | 7569 | CVCL_0493 | 1767 |

#### 3.2.1 Annotation Example

Figure 3 shows an excerpt of an annotated article on gene expression experiments.

**Fig. 3:**
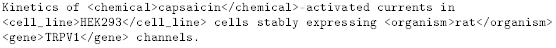
Annotation example on an OA article (PMC1266034)

### 3.3 Availability

We annotated 633,174 Open Access (OA) articles dumped on 20 July using our cell-line tagger. Those annotated OA articles will be available on Europe PMC FTP site **http://europepmc.org/ftp/oa/ner_tagging** as well as REST webservice.

## 4 Discussion

In this work, we have developed a large scale annotation and analysis system for ontologies, by exploring different technologies such as semantic web, clouding computing, etc. With this system, we have annotated and provided Open Access articles annotated with Cellosaurus-based cell line tagger on ftp site for text-mining community.

Our annotation results show that cell lines are mentioned over different sections although more often mentioned in Methods and Results sections. One surprising founding is that cell lines tagged in Results sections are less noisy then ones in Methods sections.

One application of this work is, combined with section tagger, to retrieve articles where one particular cell line mentioned in only Results sections.

As future work, we plan to extend our system adaptive and sharable using Plug and Play (P & P) annotation concept with the following features: a simple interface, dictionary P & P, semantic tagger P & P, and a feature for annotation result sharing.

## Footnotes

1 http://www.w3.org/TR/owl-features/

2 http://mallet.cs.umass.edu/

3 http://monqjfa.berlios.de/monqApiDoc/monq/programs/DictFilter.html

